# Non-canonical caspase-8 activation by cathepsin B drives anti-inflammatory human macrophage polarization

**DOI:** 10.1101/2025.08.22.671722

**Authors:** Emeline Kerreneur, Paul Chaintreuil, Chloé Delaby, Sonia Boulakirba, Maxence Bourgoin, Morgane Fajolles, Cécile Favreau, Adèle Rivault, Juba Bennour, Nathalie Droin, Jean-François Peyron, Johanna Chiche, Thomas Cluzeau, Michaël Loschi, Patrick Auberger, Guillaume Robert, Arnaud Jacquel

## Abstract

Anti-inflammatory monocyte-derived macrophages are essential to maintain tissue homeostasis but can also contribute to disease progression, notably in cancer and fibrosis. Deciphering the signaling pathways that govern their generation could therefore unlock new therapeutic opportunities. Here we uncover a previously unrecognized, non-apoptotic function of caspase-8 in driving both monocyte-to-macrophage differentiation and anti-inflammatory macrophages polarization. We identified cathepsin B as a novel upstream activator of caspase-8 activation through a non-canonical cleavage mechanism, conferring to caspase-8 an original activity profile distinct from its apoptotic role. Disruption of this cathepsin-B-caspase-8 axis, either genetically or pharmacologically, not only impairs the generation of anti-inflammatory macrophages but also reprograms these cells towards a pro-inflammatory phenotype. Our findings position the cathepsin-B-caspase-8 axis as a critical regulatory node in macrophage fate decisions and a promising target for therapeutic reprogramming of human macrophages in cancer, inflammation and fibrotic diseases.

## Introduction

Macrophages are highly versatile innate immune cells with remarkable phenotypic plasticity, shaped by their ontogeny, tissue localization and environment cues. They can be broadly classified on their origin: tissue-resident macrophages derive from erythro-myeloid progenitors (EMPs) in the fetal liver during embryogenesis^1^ whereas monocyte-derived macrophages originate from bone marrow progenitors during hematopoiesis. Hematopoietic stem cells (HSCs) give rise to common myeloid progenitors (CMPs)^2^, which differentiate into granulomonocytic progenitors (GMPs)^2^ or Monocyte-Dendritic cell progenitors (MDPs)^3^, ultimately producing mature monocytes under the influence of IL-3, GM-CSF (Granulocytes Macrophage Colony Stimulating Factor) or G-CSF (Granulocytes Colony Stimulating Factor)^3–5^. These monocytes exit the bone marrow to enter the peripheral circulation, where they constitute 5 to 10% of total blood leucocytes^6^, and further differentiate into dendritic cells upon GM-CSF and IL-4^7^ or into macrophages upon recruitment into tissues *via* diapedesis, driven by CSF-1 (Colony Stimulating Factor), CSF-2 or IL-34 signaling^8–10^.

Both tissue-resident and monocyte-derived macrophages coexist in peripheral tissues, adopting specialized phenotypes such as alveolar macrophages in the lungs, capsular macrophages in the liver and Langerhans cells in the skin^11^. While tissue-resident macrophages are long-lived and self-renewing throughout adulthood, monocyte-derived macrophages contribute to tissue maintenance during homeostasis and play key roles in inflammatory and pathological settings^11,12^. Despite their divergent origins and tissue-specific functions, macrophages share core immunological activities, including phagocytosis, antigen presentation, and the secretion of growth factors and inflammatory signaling molecules^12^.

Macrophages are also critically involved in the pathophysiology of numerous diseases. In metabolic disorders associated with obesity, they accumulate in adipose tissue and drive systemic inflammation through the release of inflammatory mediators^13^. In cancer, macrophages are instrumental in both initiation and progression of solid and hematopoietic malignancies. Tumor-associated macrophages (TAMs) and leukemia-associated macrophages (LAMs) promote tumor growth, angiogenesis and immune evasion, contributing to resistance to therapy^14,15^. Consequently, various therapeutic strategies have been developed to modulate macrophage activity in tumors, including approaches that inhibit their recruitment, deplete their population, block their activation, or reprogram them towards anti-tumoral phenotypes^6^. Understanding the molecular mechanisms that govern the differentiation and function of monocyte-derived macrophages is therefore essential to the development of novel therapeutic approaches.

*In vivo,* macrophages are continuously shaped by their microenvironment, existing along a functional continuum from pro-inflammatory to anti-inflammatory states. However, monocyte-derived macrophages are typically e*x vivo* categorized into three functional subsets. Naïve (M0-like) macrophages arise from monocyte differentiation in response to CSF-1 and can subsequently polarize into either pro-inflammatory macrophages (M1-like) or anti-inflammatory macrophages (M2-like). M1-like macrophages are induced by LPS and IFN-γ or TNF-α and IFN-γ co-stimulation and are associated with antimicrobial and anti-tumoral responses. Conversely, M2-like macrophages which arise in response to IL-4 ± IL-13, IL-6 or IL-10 support angiogenesis and tissue repair to restore homeostasis^6^.

A network of interconnected molecular pathways contributes to the survival and proper differentiation of monocytes into macrophages *ex vivo*. Specifically, CSF-1 stimulation activates the MAPK/ERK^16^, SRC kinase^17^, and PI3K/AKT pathways^18^, alongside autophagy-related mechanisms^19,20^ and the caspase cascade^8,21^. Despite these advances, the molecular mechanisms underpinning the polarization of anti-inflammatory macrophages remain incompletely understood, beyond the well-characterized role of STAT6 activation downstream of IL-4 signaling^22^. Notably, recent studies have revealed a sustained, non-apoptotic activation of caspases during anti-inflammatory macrophages polarization^23^. In particular, caspase-8^18,24,25^, caspase-3^8,18,23^ and caspase-7^23^ have been implicated in this process, though their precise respective roles and activation mechanisms remain to be elucidated.

In this study, we explore the mechanisms of non-apoptotic caspase activation, particularly that of caspase-8, during the generation of anti-inflammatory macrophages. We further evaluate the functional significance of this pathway and assess its potential as a therapeutic target for macrophage reprogramming and clinical applications.

## Materials and Methods

### Human primary monocytes purification

Human primary monocytes are purified from the peripheral blood of volunteered healthy donors with informed consent from the Etablissement Français du Sang (EFS, n°13-PP-11), the French National Agency for blood collection. First, the peripheral blood is centrifugated on a density gradient (CMSMSL01-01, Eurobio scientific, France) to separate peripheral blood mononucleated cells (PBMCs) from other blood components (plasma, neutrophils, erythrocytes). Then, PBMCs are hemolyzed (BD Pharm Lyse^TM^ Lysing Buffer, 555899, BD Biosciences, New Jersey, USA) and marked with CD14 microbeads (130-050-201, Miltenyi, Germany) to perform a positive selection using an autoMACS® Pro Separator (Miltenyi) and purify human primary monocytes.

### Human primary monocytes culture

Human primary monocytes are grown at 37°C under 5% CO_2_ in RPMI 1640 Glutamax-I (61870-044, Gibco, Massachusetts, USA) supplemented with 10% fetal bovine serum (CVFSVF06-01, Eurobio scientific) and 1% penicillin/streptomycin (15140-122, Gibco). Monocytes are then stimulated with 50ng/mL CSF-1 (130-096-493, Miltenyi) to generate naïve human primary macrophages (M0). After five days, M0 macrophages are polarized into pro-inflammatory (M1) with 100ng/mL LPS (5974-43-02, Invivogen, California, USA) + 20ng/mL IFN-γ (300-02, Peprotech, New Jersey, USA) or into anti-inflammatory macrophages (M2) with 20ng/mL IL-4 (130-094-117, Miltenyi) with or without 20ng/mL IL-13 (130-112-409, Miltenyi), 20ng/mL IL-6 (130-093-932, Miltenyi) or 20ng/mL IL-10 (78024, STEMCELL Technologies, Canada).

### Immunoblot assays

Cells are lysed for 30min at 4°C in the following lysis buffer: 50mM HEPES pH 7.4 (1560-056, Gibco), 150mM NaCl, 20mM EDTA, PhosphoSTOP (04906837001, Sigma-Aldrich, Massachusetts, USA), complete protease inhibitor (11836170001, Sigma-Aldrich) and 1% Triton X-100 (T9284, Sigma-Aldrich). Lysates are centrifugated for 15min at 16.000 g at 4°C. Supernatants are collected and dosed by spectrophotometry with Bradford solution (5000006, Bio-Rad, California, USA). An average of 50µg of proteins is diluted with the appropriate volume of PBS and Laemmli 4X (60mM Tris-HCl, 2% SDS, 10% glycerol, 0.01% bromophenol blue, 20% β-mercaptoethanol).

Samples and molecular weight markers (26619, Thermo Fisher Scientific, Massachusetts, USA) are then loaded on a polyacrylamide gel to migrate proteins in a TG-SDS solution and further transferred on a PVDF membrane (IPVH00010, Sigma-Aldrich) in a TG-20% Ethanol solution. Membranes are saturated with Blocking Buffer (Tris 10mM pH 7.4, NaCl 150mM, EDTA 1mM, Gelatin 0.5%, BSA 3%, Tween20 0.1%) for 1h at room temperature and incubated overnight at 4°C with the appropriate primary antibody. Membranes are washed three times with PBS-Tween20 0.1% and incubated with the corresponding secondary antibody for 1h30 at room temperature. Membranes are washed three more times and proteins are revelated on a photographic film with Amersham^TM^ ECL western blotting detection reagent (RPN2106, Cytiva, Massachusetts, USA).

Primary and secondary antibodies are mostly purchased from Cell Signaling Technology® (Massachusetts, USA) including Caspase-8 (9746), Caspase-7 (9492), Caspase-3 (9662), Phox (4312), Lyn (2732), HSP60 (12165), ATG7 (855S8), AIF (5318), EEA1 (3228), HRP-linked Ig mouse (7076) and HRP-linked Ig rabbit (7074). Others are from Santa Cruz Biotechnology (California, USA) including CTSL (sc32320) and LAMP2 (sc18822). The CTSB antibody is purchased from Sigma (IM27L).

### Enzymatic measurement activity

Cells are lysed for 30min at 4°C in the following lysis buffer: 50mM HEPES pH 7.4 (1560-056, Gibco), 150mM NaCl, 20mM EDTA, 0.2% Triton X-100 (T9284, Sigma-Aldrich) and 50µM PMSF. Lysates are centrifugated for 15min at 16.000 g at 4°C. Supernatants are collected and dosed by spectrophotometry with Bradford solution (5000006, Bio-rad). 10µg of lysates or recombinant proteins are deposited in quadruplicates in a 96-black-well plate with 250µM of substrates-AMC (Peptanova, Germany), 5mM of DTT and 20µM of the specific substrate inhibitor -CHO in one of the quadruplicates to remove background signals. Specific enzymatic activities are analyzed with a kinetic measure of AMC signal (every 2min, for 1h30) at an excitation wavelength of 390nm and an emission wavelength of 460nm with Biotek Synergy H1 (BioTek Instruments, Vermont, USA).

Both Ac-NKFD-AMC and Ac-KWFD-AMC substrates measure non-apoptotic caspase activities while Ac-FR-AMC, Ac-RR-AMC, Ac-IETD-AMC and Ac-DEVD-AMC substrates measure CTSB/L, CTSB, apoptotic caspase-8 and apoptotic caspase-3 activity respectively.

### Subcellular fractionation

Subcellular fractionation is performed at 4°C on PBS-washed cells with the Proteoextract® Subcellular Proteome Extraction Kit (539790, Sigma-Aldrich). After 10min of lysis of 15 x 10^6^ cells per condition with extraction buffer I and 10min centrifugation at 800g, supernatants are collected (F1: cytosolic fraction) and extraction buffer II is added to the pellet. After 30min of lysis and 10min centrifugation at 5.600g, supernatants are collected (F2: microsomal fraction). The fractioned samples are analyzed with immunoblot assays.

### Organelle isolation

Organelle isolation is performed with the lysosome isolation kit (LYSISO1, Sigma-Aldrich). After washing of a 100 x 10^6^ cells per condition with homogenization buffer made with 10mM HEPES pH 7.4 (1560-056, Gibco), 10mM KCl and 1.5mM MgCl2, cells are lysed by nitrogen cavitation, centrifugated at 1.000g for 10min at 4°C to remove nuclei and membranes and supernatants are centrifugated at 20 000g for 20min at 4°C to remove cytosol. Remaining pellets containing lysosomes and mitochondria are placed on an Optiprep® density gradient according to the manufacturer’s protocol and ultra-centrifugated at 150.000g for 4h at 4°C. Four different enriched lysosomal fractions are collected, containing lysosomes (F1 and F2), mitochondria (F3) and endosomes (F4) respectively. After centrifugation at 20.000g for 15min at 4°C, pellets are further analyzed with immunoblot assays.

### Immunofluorescence

Cells are cultured on sterilized slides within 12-well plate. After fixation with 4% PBS-FA, cells are permeabilized for 20min at −20°C with cold 70% ethanol. Cells are then saturated with PBS-BSA 8% (P06-1403500, PanBiotech, Germany) for 30min, incubated in the dark for 1h with primary antibodies and then 30min with secondary antibodies at 1/400 and 1/500 concentration respectively, previously diluted with PBS-BSA 1%. Cells are then incubated with DAPI (D9542, Sigma Aldrich) for 5 minutes and slides are mounted on glass slides with Fluoromount-G (0100-01, Southern Biotech, Alabama, USA) to be further analyzed by confocal microscopy (Nikon A1R, Japan) at 60X.

Caspase-3 and cleaved caspase-3 antibodies are purchased from Cell Signaling Technology® (9662 and 9664 respectively) and TOM20 from Santa Cruz Biotechnology (sc17764). All secondary antibodies are purchased from Invitrogen (California, USA) including Donkey anti-Rabbit IgG (H+L) Alexa Fluor™ 488 (A-21206), Goat anti-Mouse IgG2a Alexa Fluor Fluor™ 568 (A-21134) and Goat anti-Mouse IgG1 Alexa Fluor Fluor™ 647 (A-21240). Image analysis is performed using ImageJ and colocalization is evaluated by the Pearson’s correlation coefficient (r) with JACoP plugin^26^.

### Inhibitors

Bafilomycin A1 (10nM, 1334, Tocris Bioscience, UK) inhibit lysosomal acidification by targeting V-ATPase and therefore cathepsin activities. CA-074 (10 µM, S7420, Selleckchem, Texas, USA) inhibits CTSB. Emricasan (3µM, S7775, Selleckchem) is a pan-caspase inhibitor.

### Human primary macrophage transfection with siRNA

100nM siRNAs are incubated with Lipofectamine® RNAimax reagent (13778150, Invitrogen) in a 1:2 ratio in Opti-MEM™ I (31985070, Gibco) for 5min before transfection of macrophages at day 5 of differentiation for 48h or 72h with 20ng/mL of IL-4. siLuc (Custom select 4390829, Invitrogen) is used as a control of transfection. siCaspase-8 (On target plus smart pool #L-003466-00-0020, Dharmacon, Colorado, USA) is a pool of 4 target sequences. Other siRNAs are purchased from Invitrogen with the following references: siCaspase-7 (C7HSS101381), siCaspase-3 (C3HSS101372), siCTSB (CTSBHSS102477), siCTSD (CTSDHSS102478), siCTSL (CTSLHSS102494), siCTSS (CTSSHSS102502).

### *In vitro* cleavage of Caspase-8

Recombinant caspase-8 (50ng, Enzo Life Sciences, New York, USA) with or without recombinant active CTSB (400ng, Sigma-Aldrich) and CA-074 (10µM, S7420, Selleckchem) are incubated in acidic medium (NaH2PO4 250mM, Na2HPO4 125mM, EDTA 2mM, DTT 10µM) for 24h at 37°C. *In vitro* cleavage of caspase-8 is evaluated with immunoblot assays.

### RNA extraction, reverse-transcription and real-time quantitative polymerase chain reaction

RNA is extracted from 5 x 10^6^ cells per condition with RNeasy® Mini kit (74106, Qiagen, Germany) according to manufacturer’s protocol and concentration is measured with Nanodrop (Thermo Fisher Scientific) at 260nm. cDNA is produced from 1000ng of RNA with Random primers (C118A), dNTP (100µM, U1515), rec RNAsin® (N251A), AMV RT (M510F) and AMV RT 5X Buffer (M515A) from Promega (Wisconsin, USA), following standard protocols. Real-time polymerase chain reaction (PCR) is performed with PowerUp™ SYBR™ Green Master Mix protocol (A25742, Applied Biosystems, California, USA), with 500nM of each appropriate primers, available upon request. L32 is used as a control of endogenous expression.

### Flow cytometry

Cells are labeled with 1µL of the appropriate antibody diluted with 50µL of MACSQuant® Running Buffer (130-092-747, Miltenyi) for 10min at 4°C and further fixed with PBS-PFA 4% (15710, Electron Microscopy Sciences, Pennsylvania, USA). Fluorescence is measured on a MACSQuant10 analyzer (Miltenyi). Antibodies are purchased from Miltenyi including CD200R (130-111-291), CD209 (130-120-729) and CD86 (130-116-159).

### ELISA

48h (pharmacologic inhibition) or 72h (genetic inhibition) macrophage supernatants are diluted with commercial diluent and further loaded into the commercial plate. Concentration is analyzed with the Ella Automated Immunoassay System (Bio-Techne, Minnesota, USA). Dilution factors are the following: 250 for CCL18 and CCL26 for pharmacologic inhibition samples, 100 for CCL18 and CCL26 for genetic inhibition samples, 2 for TNF-α and CCL20 for both pharmacologic and genetic inhibition samples. Results are further rationalized to the number of secreting cells evaluated by flow cytometry and to the control condition.

### Whole-transcriptome RNA-sequencing

The RNA integrity (RNA Integrity Score ≥ 7.0) was checked on the Agilent Fragment Analyzer (Agilent Technologies, California, USA) and quantity was determined using Nanodrop. SureSelect Automated Strand Specific RNA Library Preparation Kit was used according to manufacturer’s instructions with the Bravo Platform (Agilent Technologies). Briefly, 200ng of total RNA per sample was used for poly-A mRNA selection using oligo(dT) beads and subjected to thermal mRNA fragmentation. The fragmented mRNA samples were subjected to cDNA synthesis and were further converted into double stranded DNA using the reagents supplied in the kit, and the resulting dsDNA was used for library preparation. The final libraries were indexed, purified, pooled together in equal concentrations and subjected to paired-end sequencing (2x100 bp) on Novaseq-6000 sequencer (Illumina, California, USA) at Gustave Roussy Institute.

### RNA-sequencing analysis

Raw sequencing reads of whole-transcriptome RNA-sequencing in FASTQ format were processed through a standard RNA-sequencing analysis pipeline. Quality control was carried out using FastQC (v0.11.9)^27^ and summarized with MultiQC (v1.12)^28^. Adapter sequences were trimmed using Cutadapt (v3.5) and only high-quality reads were retained for downstream processing^29^. Cleaned reads were then aligned to the Homo sapiens reference genome (GRCh38, GENCODE v47) using STAR aligner with default parameters (v.2.7.10b)^30^. Gene-level quantification was performed using featureCounts (v2.0.3) generating a raw count matrix for all samples^31^. Subsequent analyses were conducted within the R environment (v4.4.3) using Bioconductor (v3.19). Sample-level quality control included dimensionality reduction *via* Principal Component Analysis (PCA) to assess potential batch effects and the global similarity between biological replicates. Differential expression analysis (DEA) was carried out using DESeq2 algorithm (v1.44.0)^32^. Differentially expressed genes (DEGs) were identified using a Benjamin-Hochberg ajusted p-value < 0.05 and an absolute log2 fold changes ≥ 1. Enrichment analysis was then performed using clusterProfiler (v4.14.4) with the Gene Ontology (GO) Biological Process database^33^. Only statistically significant terms (adjusted p-value < 0.05) were retained for interpretation. All visualizations were generated in R using ggplot2 (v3.5.1)^34^ and related tidyverse packages (v1.3.1)^35^ while heatmaps were constructed with ComplexHeatmap (v2.20.0)^36^.

### Statistical analysis

Statistical analysis is performed using GraphPad Prism 9.4.0 software on at least 3 independent experiments with unpaired two-tailed Student’s t-tests for 2-conditions experiments or ordinary one-way ANOVA for the others. Results are expressed as mean ± SEM. * p < 0.05 ; ** p < 0.01 ; *** p < 0.001 ; **** p < 0.0001.

## Results

### Non-apoptotic caspases are activated during monocyte-to-macrophages differentiation and anti-inflammatory macrophages polarization

We previously demonstrated that *ex vivo* exposure of primary monocytes to CSF-1 induces the non-apoptotic activation of caspase-8 *via* the assembly of a multiprotein complex comprising caspase-8, FADD, RIPK1, and FLIP proteins^25^. This atypical activation observed three days after CSF-1 treatment, resulted in non-canonical cleavages of caspase-8, caspase-7, caspase-3 and their substrates p47 Phox and Lyn, generating fragments of 34 kDa, 30 kDa, 26 kDa, 40 kDa and 51 kDa, respectively (Fig. 1A). In contrast, classical apoptotic cleavage patterns were only observed in unstimulated monocytes (day 0), likely reflecting residual apoptosis induced by cell sorting, and were no longer detectable following CSF-1 stimulation (Fig. 1A). To assess the persistence of this unconventional caspase activation during the macrophages functional polarization, we differentiated CSF-1-treated monocytes into either pro-(M1) or anti-inflammatory (M2) macrophages by stimulating unpolarized macrophages (M0) with LPS + IFN-γ or IL-4, respectively. Strikingly, non-canonical cleavage of caspases and their substrates persisted in M2 macrophages but was completely abrogated in M1 macrophages (Fig. 1B), underscoring a selective activation mechanism associated with CSF-1-driven and M2-like macrophages. Proteomic characterization of more of a dozen caspase substrates^37–39^ in CSF-1 treated monocytes revealed non-canonical cleavage sites preferentially involving NxxD or KxxD motifs (Supplementary Fig. 1A). To selectively monitor this non-apoptotic caspase activity during macrophage activation/polarization, we developed two novel fluorescent synthetic substrates, Ac-NKFD-AMC and Ac-KWFD-AMC. We demonstrated that these substrates were specifically cleaved in M0 and IL-4-polarized macrophages, but not in M1 macrophages or apoptotic monocytes (Fig. 1C and Supplementary Fig. 1B), consistent with Western blot analyses. In contrast, standard caspase activity assays using Ac-IETD-AMC and Ac-DEVD-AMC failed to distinguish between polarization states (Supplementary Fig. 1C and 1D). Notably, recombinant active caspase-8 or caspase-3 did not cleave Ac-NKFD-AMC or Ac-KWFD-AMC, whereas they efficiently processed the canonical substrates Ac-IETD-AMC and Ac-DEVD-AMC (Supplementary Fig. 1E and 1F). Given the plasticity of macrophages, we next examined whether this caspase activation pattern was reversible. We thus generated M1- and M2-like macrophages *ex vivo* and repolarized them for two days using IL-4 or LPS + IFN-γ respectively (Fig. 1D and 1E). IL-4 reinduced both caspase cleavage (Fig. 1D) and non-apoptotic caspase activity (Fig. 1E), whereas LPS + IFN-γ abrogated it. Finally, we showed that this non-apoptotic activation of caspases also occurred in response to IL-4 + IL-13, IL-6, and IL-10 stimulation, highlighting its broader relevance in the generation of all M2-like macrophages subsets (Supplementary Fig. 1G and 1H).

**Fig. 1.**
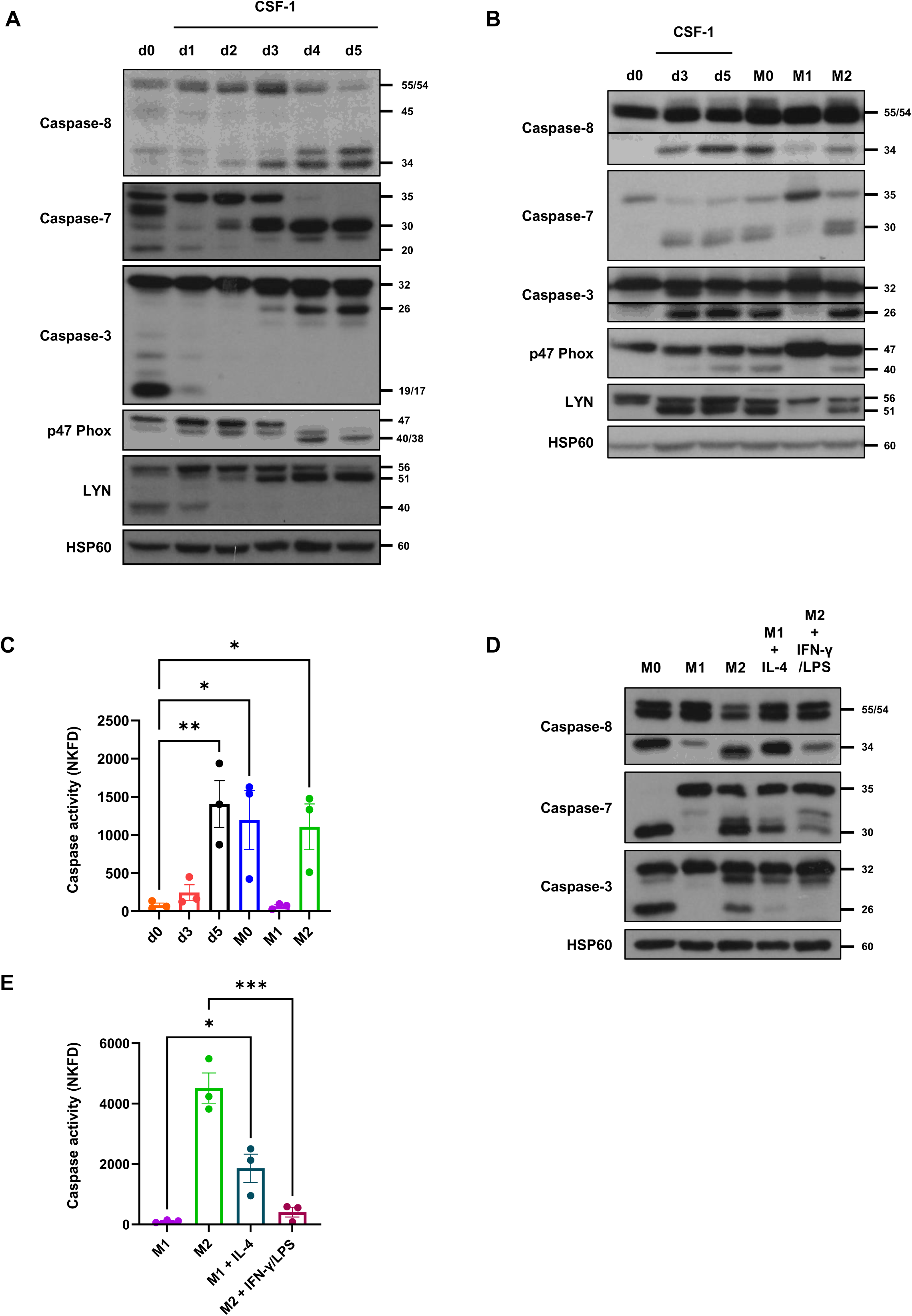
Non-apoptotic caspase activation during monocyte-to-macrophage differentiation and macrophage polarization. Human monocytes (d0) are differentiated with CSF-1 for 5 days and then treated with CSF-1 (M0) or polarized with IFN-γ + LPS (M1) or IL-4 (M2) for 2 days. **a-b** Caspase-8, -7 and -3, p47 Phox and Lyn expression analysis by immunoblotting. HSP60 is used as a loading control. **c** Enzymatic measurement of non-apoptotic caspase activities using a specific fluorescent peptide (Ac-NKFD-AMC). Results are expressed as A.U/min/mg proteins and represent the mean ± SEM of 3 independent experiments performed in triplicates. After 2 days of polarization, pro-inflammatory macrophages are exposed to IL-4 and anti-inflammatory macrophages are submitted to IFN-γ + LPS for 2 days. **d** Caspase-8, -7 and -3 expression analysis by immunoblotting. HSP60 is used as a loading control. **e** Enzymatic measurement of non-apoptotic caspase activities using a specific fluorescent peptide (Ac-NKFD-AMC). Results are expressed as A.U/min/mg proteins and represent the mean ± SEM of 3 independent experiments performed in triplicates. * p < 0.05, ** p < 0.01 and *** p < 0.001 according to an ordinary one-way ANOVA.

### Non-apoptotic cleavage fragments of caspases are specifically located in the mitochondrial compartment in unpolarized and anti-inflammatory macrophages

To further investigate the specificity of these non-apoptotic caspase fragments, we examined their subcellular localization as apoptotic fragments are typically confined to the cytosol^40^. Subcellular fractionation of differentiating and polarized macrophages revealed that non-apoptotic fragments of caspases are predominantly enriched in the microsomal fraction (Fig. 2A), and more specifically within the mitochondrial / endosomal compartment (Fig. 2B). Immunofluorescence analysis confirmed that the non-apoptotic fragment of caspase-3 in both unpolarized and anti-inflammatory macrophages is restricted to mitochondria (Fig. 2C), colocalizing with TOM20, a marker of the outer mitochondrial membrane^41^. This atypical localization highlights the distinct and potentially specific roles of non-apoptotic caspases during monocyte-to-macrophages differentiation and M2-like polarization.

**Fig. 2.**
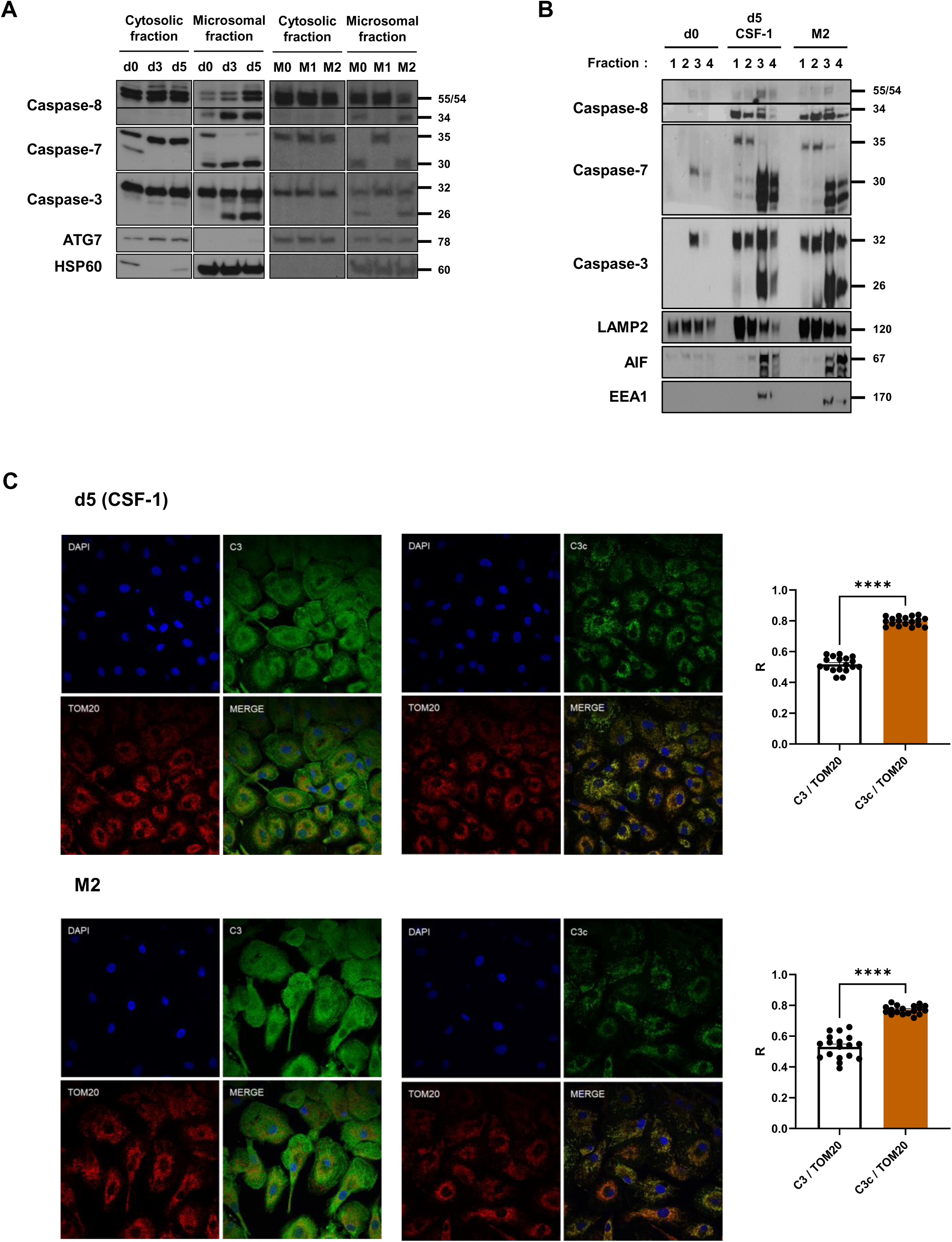
Non-apoptotic cleavage fragments of caspase localization in unpolarized and anti-inflammatory macrophages. Human monocytes (d0) are differentiated with CSF-1 for 5 days and then treated with CSF-1 (M0) or polarized with IFN-γ + LPS (M1) or IL-4 (M2) for 2 days. **a** Caspase-8, -7 and -3 localization analysis by immunoblotting after subcellular fractionation of cytosolic and microsomal compartments. ATG7 and HSP60 are used as controls, enriched in cytosolic or microsomal fractions respectively. **b** Caspase-8, -7 and -3 localization analysis by immunoblotting after organelle isolation and lysosomal enrichment. LAMP2, AIF and EEA1 are used as controls, accumulated in lysosomes (1+2), mitochondria (3) and endosomes (4) respectively. **c** Colocalization analysis of caspase-3 and its non-apoptotic fragment with TOM20, a mitochondrial marker, by immunofluorescence staining. Results are expressed with Pearson’s coefficient (R) and represent the mean ± SEM of 3 independent experiments performed in sextuplicate. **** p < 0.0001 according to a two-tailed unpaired Student’s t test.

### Non-apoptotic caspase-7 and caspase-3 cleavages rely on caspase-8 activation while Cathepsin B mediates the non-apoptotic caspase-8 cleavage during monocyte-to-macrophages differentiation and anti-inflammatory macrophages polarization

Caspase-7 and caspase-3 are executioner caspases, typically activated by initiator caspases such as caspase-8 during apoptosis^40^. To explore how these caspases are activated in a non-apoptotic context during monocyte-to-macrophage differentiation and anti-inflammatory macrophages polarization, we treated cells with Emricasan, a pan-caspase inhibitor^42^. Immunoblot analysis revealed that the cleavage of caspase-7 and caspase-3 but not caspase-8 was abolished by Emricasan treatment in both differentiating and anti-inflammatory macrophages (Fig. 3A). Of note, targeted silencing of caspase-8 using siRNA disrupted the generation of non-apoptotic fragments of caspase-7 and caspase-3 in both cell types (Fig. 3B). Collectively, these results establish that non-apoptotic activation of caspase-7 and caspase-3 is mediated by caspase-8 during macrophage differentiation and M2-like polarization.

**Fig. 3.**
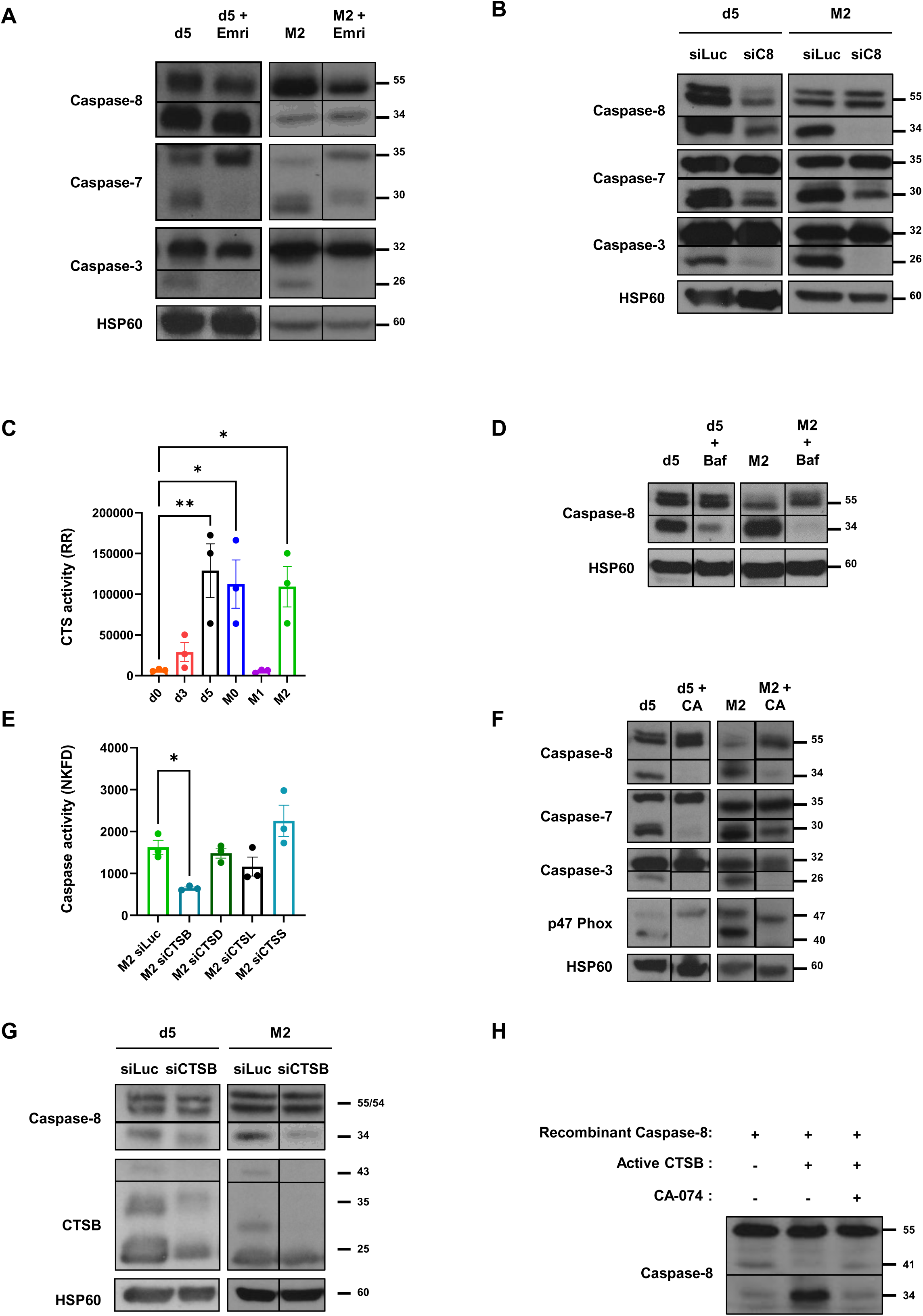
Non-apoptotic cleavage of caspase-7 and -3 by caspase-8 and of caspase-8 by CTSB during monocyte-to-macrophage differentiation and macrophage polarization. Differentiating and anti-inflammatory macrophages are treated with Emricasan at d4. M2 macrophages are polarized during two days with IL-4 with or without Emricasan. Caspase-8, -7 and -3 expression analysis by immunoblotting. HSP60 is used as a loading control. **b** Differentiating and anti-inflammatory macrophages are transfected with siRNA targeting Luciferase (control) or Caspase-8 for 3 days. Caspase-8, -7 and -3 expression analysis by immunoblotting. HSP60 is used as a loading control. **c** Monocytes are differentiated with CSF-1 for 5 days and treated with CSF-1 (M0), IFN-γ + LPS (M1) or IL-4 (M2) for 2 days. CTSB+L enzymatic activity is measured using Ac-RR-AMC substrate. Results are expressed as A.U/min/mg proteins and represent the mean ± SEM of 3 independent experiments performed in triplicates. **d** Differentiating and anti-inflammatory macrophages are treated with Bafilomycin at d4. M2 macrophages are polarized during two days with IL-4 with or without Bafilomycin. Caspase-8 expression analysis by immunoblotting. HSP60 is used as a loading control. **e** Anti-inflammatory polarizing macrophages are transfected with siRNA targeting Luciferase, CTSB, D, L or S for 2 days. Non-apoptotic caspase enzymatic activity is measured using Ac-NKFD-AMC. Results are expressed as A.U/min/mg proteins and represent the mean ± SEM of 3 independent experiments performed in triplicates. **f** Differentiating and anti-inflammatory macrophages are treated with CA-074 at d4. M2 macrophages are polarized during two days with IL-4 with or without CA-074. Caspase-8, -7, -3 and p47 Phox expression analysis by immunoblotting. HSP60 is used as a loading control. **g** Differentiating and anti-inflammatory polarizing macrophages are transfected with siRNA directed against Luciferase or CTSB for 3 days. Caspase-8 and CTSB expression analysis by immunoblotting. HSP60 is used as a loading control. **h** Recombinant Caspase-8 is incubated *in vitro* with active CTSB with or without CA-074 in an acidic medium for 24H at 37°C. Caspase-8 analysis by immunoblotting. * p < 0.05 and ** p < 0.01 according to an ordinary one-way ANOVA.

Since caspase-8 activation in this context does not depend on caspases activity and given that cathepsins, particularly cathepsin D (CTSD), have been implicated in caspase-8 activation during apoptosis^43,44^, we investigated the role of cathepsins during monocyte-to-macrophage differentiation and polarization. Using the fluorescent synthetic cathepsin substrates Ac-RR-AMC and Ac-FR-AMC, we detected a robust cathepsin activity in unpolarized and anti-inflammatory macrophages, but not in pro-inflammatory macrophages nor apoptotic monocytes (Fig. 3C and Supplementary Fig. 2A). Given that cathepsins reside in lysosomes^45^, we next inhibited lysosomal acidification by exposing unpolarized and anti-inflammatory macrophages to Bafilomycin A1, a specific inhibitor of the v-ATPase proton pump^46^. Under these conditions, the generation of the non-apoptotic caspase-8 fragment was strongly suppressed in both unpolarized and anti-inflammatory macrophages (Fig. 3D). Furthermore, non-apoptotic caspase activity was selectively impaired upon genetic inhibition of cathepsin B (CTSB) in polarized anti-inflammatory macrophages (Fig. 3E and Supplementary Fig. 2B). Both genetic (siRNA) and pharmacological (CA-074, a CTSB inhibitor) inhibition of CTSB disrupted the formation of non-apoptotic cleavage fragments of caspase-8, caspase-7, caspase-3 and p47 Phox during monocyte-to-macrophages differentiation and anti-inflammatory polarization (Fig. 3F and 3G). Finally, *in vitro* incubation of recombinant caspase-8 with active CTSB produced a 34kDa fragment identical to that observed in anti-inflammatory macrophages (Fig. 3H). Collectively, these results demonstrate that the non-apoptotic cleavage of caspase-8 is CTSB-dependent and occurs during both monocyte-to-macrophages differentiation and M2-like macrophage polarization.

### Pharmacological inhibition of CTSB and caspases in unpolarized macrophages prevents anti-inflammatory macrophages generation

We next investigated the functional role of the CTSB / non-apoptotic caspase pathway in the anti-inflammatory polarization of M2-like macrophages. In this end, unpolarized macrophages were pre-treated for 24 hours with either CA-074 or Emricasan, followed by 48 hours of stimulation with IL-4 in continued presence of the inhibitors. We first confirmed that CTSB inhibition effectively impaired both cathepsin and non-apoptotic caspases activities, whereas caspase inhibition selectively reduced non-apoptotic caspase activities (Supplementary Fig. 3A and 3B). Importantly, the inhibition of either CTSB or caspases significantly reduced the surface expression of the anti-inflammatory markers CD200R, a type I membrane glycoprotein, and CD209, a C-type lectin receptor (Fig. 4A to 4D). Both molecules play key roles in modulating immune responses and inflammation^6^. This reduction in surface marker expression correlates with the dampened expression of the CCL17 and CCL18 chemokines, indicating that pharmacological blockade of CTSB or caspases disrupts the acquisition of an anti-inflammatory macrophage phenotype (Fig. 4E and 4F).

**Fig. 4.**
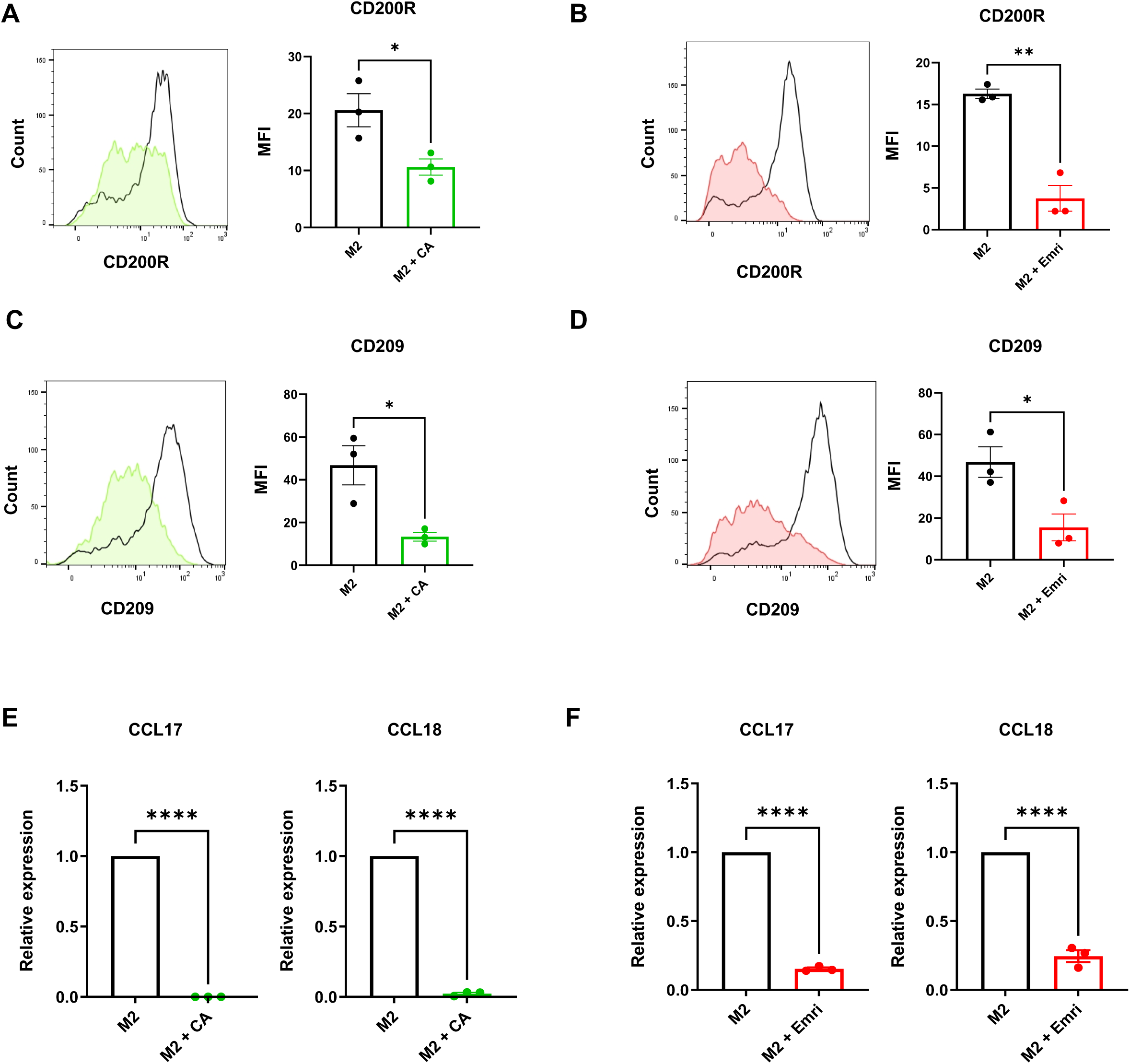
Macrophage anti-inflammatory polarization under pharmacological inhibition of CTSB and caspases. Human monocytes are differentiated with CSF-1 for 5 days. Unpolarized macrophages are treated with Emricasan or CA-074 at d4 and further polarized during two days with IL-4, in presence of the inhibitors. **a-d** Cytometry analysis of CD200R and CD209 markers expression. Results are expressed as MFI (Mean Fluorescence Intensity) and represent the mean ± SEM of 3 independent experiments. **e-f** RT-qPCR analysis of CCL17 and CCL18 gene expression. Results are expressed as relative expression and represent the mean ± SEM of 3 independent experiments. * p < 0.05, ** p < 0.01 and **** p < 0.0001 according to a two-tailed unpaired Student’s t test.

### Pharmacological inhibition of CTSB and caspases in anti-inflammatory macrophages trigger phenotypical and functional reprogramming towards pro-inflammatory macrophages

Having identified CTSB and non-apoptotic caspases as key regulators of anti-inflammatory macrophages polarization, we next investigated whether targeting this pathway could also reverse the phenotype of already polarized M2-like macrophages. To this end, macrophages were first stimulated with IL-4 for two days to induce anti-inflammatory polarization, followed by treatment with either CA-074 or Emricasan for an additional two days. We checked that CA-074 efficiently abrogated both CTSB and caspase activities, while Emricasan selectively inhibited caspase activity under the same conditions (Supplementary Fig. 4A and 4B). Strikingly, pharmacological inhibition of CTSB or caspases led to a significant reduction of the anti-inflammatory markers CD200R and CD209 (Fig. 5A to 5D). Notably, the expression of CD86, a crucial pro-inflammatory co-stimulatory molecule, was specifically increased by Emricasan (Fig. 5E and 5F). This correlated with the decreased expression of the anti-inflammatory chemokines CCL18 and CCL13 (Fig. 5G and 5H) and a concomitant increased in IL1-α and CCL20 expression, exclusively when caspases were inhibited (Fig. 5I and 5J). Thus, while CTSB targeting dampened anti-inflammatory polarization of M2 macrophages, caspase inhibition goes further by actively reprogramming them phenotypically towards a pro-inflammatory status. To confirm this switch, we assessed the functional abilities of macrophages using ELISA assays. We firstly evidenced a lower secretion of anti-inflammatory chemokines CCL18 and CCL26 by anti-inflammatory macrophages treated with CA-074 and Emricasan (Fig. 5K and 5L). Interestingly, both CTSB and caspase inhibition increased TNF-α secretion, whereas CCL20 secretion was specifically induced by Emricasan (Fig. 5M and 5N). Additionally, lactate secretion, a hallmark of anaerobic glycolysis and a metabolic feature of pro-inflammatory macrophages^47^, was also elevated upon CTSB or caspase inhibition (Supplementary Fig. 4C and 4D). Lastly, the phagocytic potential of anti-inflammatory macrophages to engulf beads coated with particles of E. Coli was significantly reduced following CTSB and caspase inhibition (Supplementary Fig. 4E and 4F). In conclusion, pharmacological inhibition of CTSB or caspases in anti-inflammatory macrophages reprograms them towards pro-inflammatory macrophages.

**Fig. 5.**
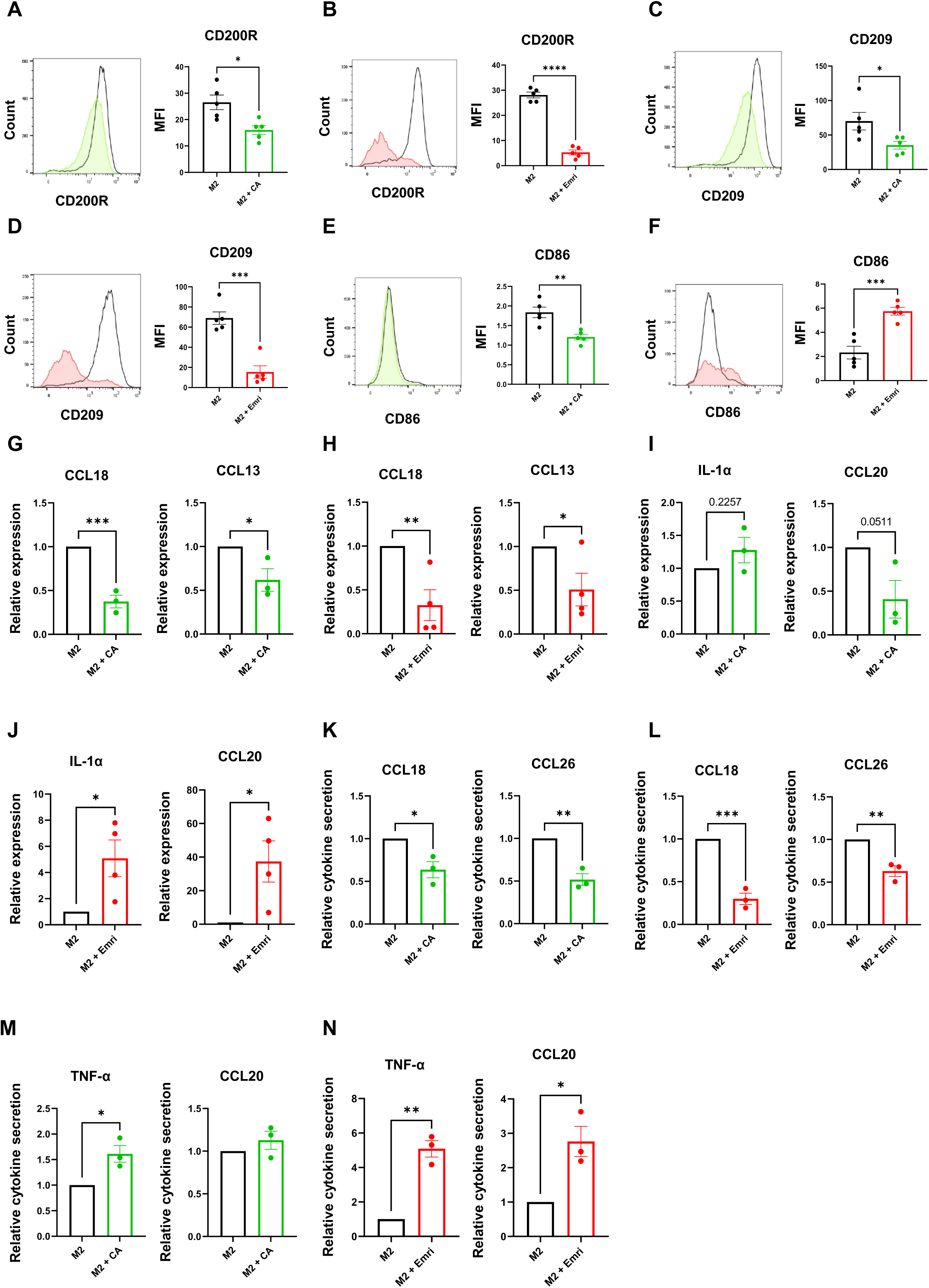
Anti-inflammatory macrophage reprogramming under pharmacological inhibition of CTSB and caspases. Human monocytes are differentiated in macrophages with CSF-1 for 5 days and polarized into anti-inflammatory macrophages during 2 days with IL-4 and further treated with Emricasan or CA-074 for 2 days. **a-f** Flow cytometry analysis of CD200R, CD209 and CD86 markers expression. Results are expressed as MFI (Mean Fluorescence Intensity) and represent the mean ± SEM of 5 independent experiments. **g-j** RT-qPCR analysis of CCL18, CCL13, IL-1α and CCL20 gene expression. Results are expressed as relative expression and represent the mean ± SEM of 3 (**g, i**) or 4 (**h, j**) independent experiments. **k-n** Secretion analysis of CCL18, CCL26, TNF-α and CCL20 by ELISA assay. Results are expressed as relative cytokine secretion (pg/mL/10^6^ cells) and represent the mean ± SEM of 3 independent experiments. ns : p > 0.05 ; * p < 0.05 ; ** p < 0.01 ; *** p < 0.001 and **** p < 0.0001 according to a two-tailed unpaired Student’s t test.

### Genetic inhibition of CTSB or caspase-8 reprogram anti-inflammatory macrophages into pro-inflammatory macrophages

To confirm the reprogramming potential of CTSB and caspase inhibition on anti-inflammatory macrophages, we genetically silenced key components of the pathway: CTSB, caspase-8, caspase-7, caspase-3 or luciferase (transfection control) and CTSL (as negative control). IL-4-treated macrophages were transfected during three days with siRNAs. Target silencing was verified by RT-qPCR (Supplementary Fig. 5A) and corresponding decrease in CTS and non-apoptotic caspase activities were confirmed (Supplementary Fig. 5B). Surface expression of anti-inflammatory markers CD200R and CD209 was significantly reduced upon knockdown of CTSB, caspase-8, caspase-7 and caspase-3 (Fig. 6A and 6B). Interestingly, CD86 was upregulated specifically when caspase-8 was silenced (Fig. 6C). Of note, CTSB or caspase silencing correlated with decreased expression of CCL18 and CCL13, while CTSL inhibition was ineffective (Fig. 6D). More interestingly, in the same conditions, we highlighted the overexpression of IL1-α and a specific overexpression of CCL20 when caspase-8 is inhibited, traducing a better efficiency of its reprogramming potential on anti-inflammatory macrophages compared to caspase-7, caspase-3 and CTSB (Fig. 6E). Given these findings, we focused on CTSB or caspase-8 knockdown for functional assays. Both conditions resulted in a decreased secretion of anti-inflammatory chemokines CCL18 and CCL26 (Fig. 6F). However, only caspase-8 silencing increased TNF-α and CCL20 secretion (Fig. 6G). Finally, we evidenced that the phagocytic potential of anti-inflammatory macrophages is decreased by CTSB or caspase-8 silencing (Supplementary Fig. 5C). Collectively, these results demonstrate that the genetic inhibition of CTSB, caspase-7, caspase-3 and most prominently caspase-8 reprograms anti-inflammatory macrophages towards a pro-inflammatory phenotype.

**Fig. 6.**
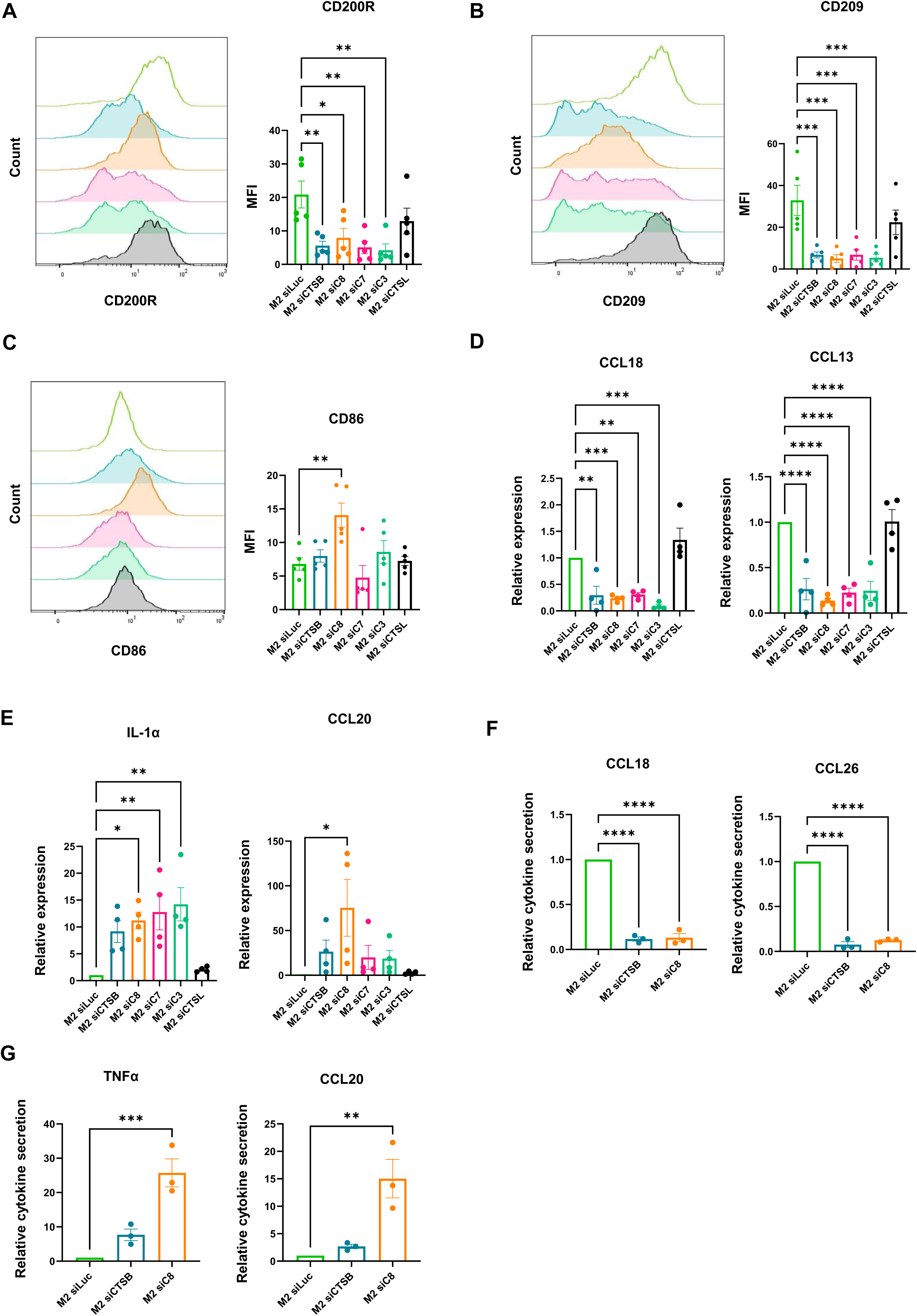
Anti-inflammatory macrophage phenotypical and functional reprogramming under genetic inhibition of CTSB and caspase-8. Anti-inflammatory macrophages are transfected with siRNA directed against Luciferase (used as a control of transfection), caspase-8, -7, -3, CTSB or L during 2 (**d-e**) or 3 days (**a-c**, **f-g**). **a-c** Flow cytometry analysis of CD200R, CD209 and CD86 marker expression. Results are expressed as MFI (Mean Fluorescence Intensity) and represent the mean ± SEM of 5 independent experiments. **d-e** RT-qPCR analysis of CCL18, CCL13, IL-1α and CCL20 gene expression. Results are expressed as relative expression and represent the mean ± SEM of 4 independent experiments. **f-g** Secretion analysis of CCL18, CCL26, TNF-α and CCL20 by ELISA assay. Results are expressed as relative cytokine secretion (pg/mL/10^6^ cells) and represent the mean ± SEM of 3 independent experiments. ns : p > 0.05 ; * p < 0.05 ; ** p < 0.01, *** p < 0.001 and **** p < 0.0001 according to an ordinary one-way ANOVA.

### Transcriptomic reprogramming of anti-inflammatory macrophages upon genetic inhibition of CTSB and caspase-8

To better assess the global transcriptional reprogramming induced by caspase-8 or CTSB inhibition, we performed a transcriptomic profiling. After validating caspase-8 or CTSB extinction (Supplementary Fig. 6A), principal component analysis revealed clear clustering of experimental groups (Fig. 7A and Supplementary Fig. 6B). Caspase-8 inhibition significantly altered the expression of 3211 genes (absolute log2 fold change ≥ 1, p-value < 0.05), with 1615 upregulated and 1596 downregulated (Fig. 7B). Gene ontology enrichment analysis of the top 20 biological processes revealed a strong activation of several pro-inflammatory pathways including “response to cytokine”, “response to bacterium”, “inflammatory response” and “leukocyte activation” (Fig. 7C). A heatmap focusing on differentially expressed genes (absolute log2 fold change > 1.5) enriched in at least two of these GO categories revealed a decreased expression of anti-inflammatory chemokines, such as CCL23 and CCL24, and increased expression of pro-inflammatory chemokines, including CCL4 and CCL7 (Fig. 7D). These transcriptomic changes were further validated by RT-qPCR (Fig. 7E). In contrast, CTSB silencing resulted in a more limited transcriptomic response, with only 218 significantly dysregulated genes (96 up-regulated, 122 down-regulated) (Supplementary Fig. 6C). Despite this, five GO categories were found commonly enriched, including “response to cytokine” and “inflammatory response” in both CTSB and caspase-8 silencing (Supplementary Fig. 6D). Altogether, these transcriptomic data emphasize the pivotal role of the CTSB / caspase-8 axis in orchestrating the phenotypic and functional reprogramming of anti-inflammatory macrophages towards a pro-inflammatory identity.

**Fig. 7.**
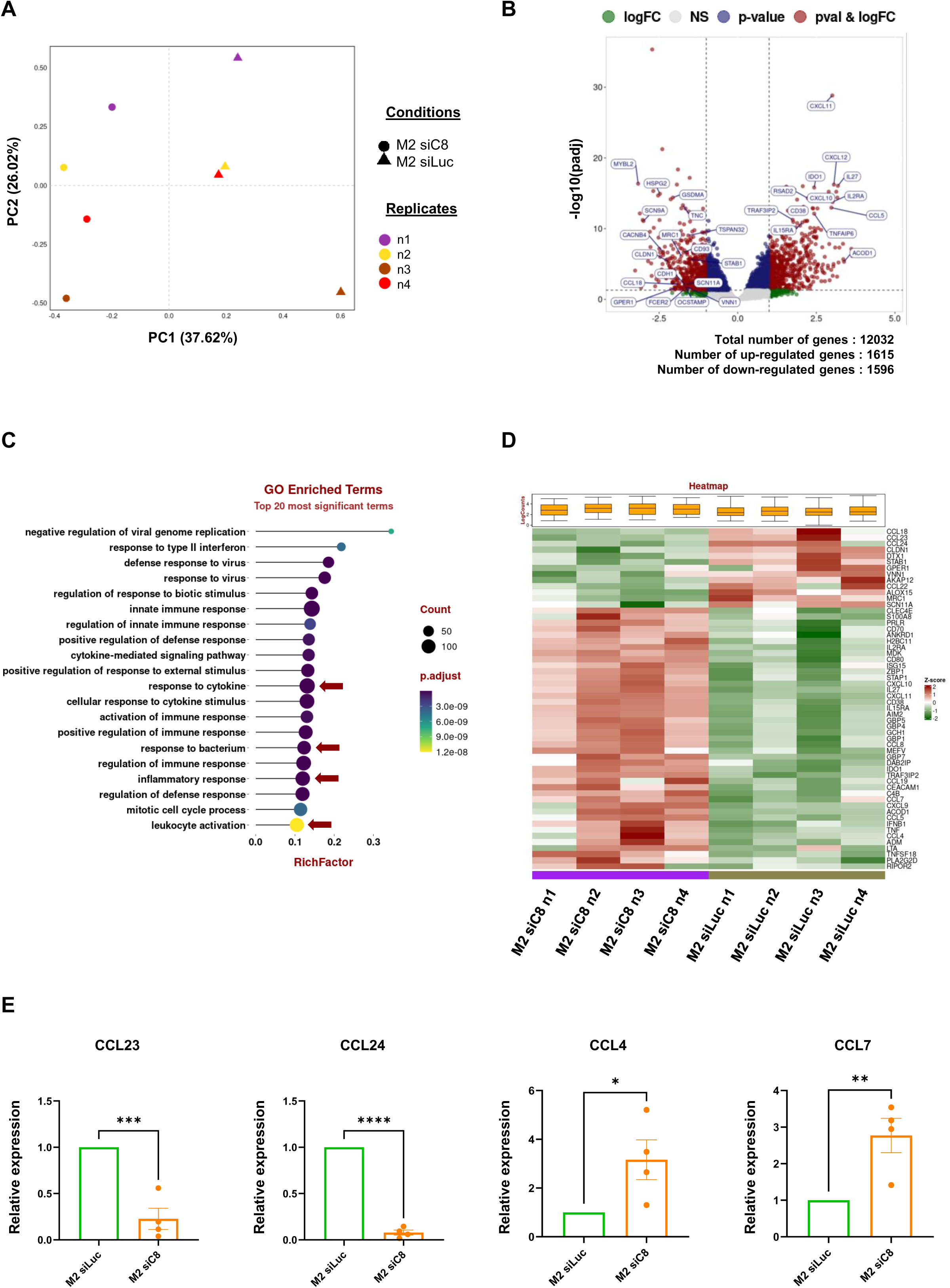
Transcriptomic reprogramming of anti-inflammatory macrophages under genetic inhibition of caspase-8. Anti-inflammatory macrophages are transfected with siRNA directed against Luciferase (M2 siLuc), used as a control of transfection, or Caspase-8 (M2 siC8) for 2 days. **a** Unsupervised 2D projection of the eight transcriptome samples principal component analysis. **b** Volcano plot representation of significant dysregulated genes between M2 siC8 compared to M2 siLuc. **c** Dot plot representation of the top 20 biological processes from gene ontology analysis. **d** Heatmap of a selected panel of genes, significantly dysregulated and with absolute log2 fold change > 1.5 in at least two GO biological processes among *response to cytokine*, *response to bacterium*, *inflammatory response* and *leukocyte activation*. **e** RT-qPCR analysis of CCL23, CCL24, CCL4 and CCL7 gene expression. Results are expressed as relative expression and represent the mean ± SEM of 4 independent experiments. ns : p > 0.05 ; * p < 0.05 ; ** p < 0.01 and *** p < 0.001 according to a two-tailed unpaired Student’s t test.

## Discussion

In the present study, we uncover the crucial role of CTSB and non-apoptotic caspases in both the *ex vivo* differentiation of human primary monocytes into macrophages and the polarization of anti-inflammatory macrophages. We previously reported that caspase-8 is activated in a non-apoptotic manner through a complex involving FADD, RIPK1 and FLIP, upon CSF-1R stimulation by CSF-1^25^ or IL-34^6,10^. Here we demonstrate that CTSB mediates the initial activation of this non-apoptotic caspase cascade in response to CSF-1, without triggering cell death. Notably, we show that the CTSB-caspase-8 axis is also activated during anti-inflammatory macrophages polarization following IL-4, IL-6 or IL-10 stimulation. In contrast, pro-inflammatory polarization with LPS and IFN-γ abolish non-apoptotic caspase activities. This observation led us to investigate whether non-apoptotic caspase activity is initiated during differentiation and sustained through anti-inflammatory polarization, or whether it can be independently re-induced. We found that IL-4 stimulation alone of polarized pro-inflammatory macrophages was sufficient to reactivate non-apoptotic caspase activity. This is likely the result of Akt activation, previously implicated in caspase activation during monocyte-to-macrophage differentiation^18^ and also induced upon IL-4 stimulation^48^. Conversely, stimulation of already polarized anti-inflammatory macrophages with LPS + IFN-γ abolished non-apoptotic caspase activity. One plausible explanation involves natural CTSB inhibitors such as cystatin C which are upregulated in M1 macrophages and for which a decrease in secretion has previously been observed in IFN-γ-treated mouse peritoneal macrophages^49^. Another likely mechanism could be the higher lysosomal pH of M1 compared to M2 macrophages, that may impede CTSB activation. Together, these findings highlight the complexity of macrophage polarization and identify CTSB as an initiator of a non-apoptotic caspase cascade in anti-inflammatory macrophages.

We further demonstrate that activation of the non-apoptotic caspase cascade during differentiation and polarization lead to the original cleavage of multiple protein substrates including nucleophosmin (NPM)^37,39^, p47 Phox^38^ (Fig. 1A), Lyn (Fig. 1A) and Beclin-1 (data not shown). Detailed analysis of their cleavage sites revealed two conserved canonical cleavage motif “KxxD” or “NxxD”, that were systemically found in dozens of substrates so far identified^37–39^. Based on these sequences, we designed two selective fluorogenic substrates, Ac-KWFD-AMC and Ac-NKFD-AMC, derived from p47 Phox and Beclin-1 respectively. These substrates were not cleaved by active apoptotic caspase-8 or caspase-3 *in vitro*, allowing for the specific detection of non-apoptotic caspase activity in *ex vivo* monocyte-derived macrophages. The functional role of two of the identified non-apoptotic caspase substrates has been previously assessed in CSF-1 treated monocytes. The non-apoptotic cleavage of NPM was shown to suppress macrophage phagocytosis and motility^39^, while the caspase-7-dependent cleavage of p47 Phox, promotes NOX2 complex activation and the subsequent production of cytosolic reactive oxygen species^38^. In that study, Solier et al. localized active caspase-3 and 7 to the outer mitochondrial membrane during macrophage differentiation. We confirm this subcellular localization by highlighting the co-localization of cleaved caspase-3 with TOM20, a mitochondrial outer membrane marker, in both differentiated and anti-inflammatory macrophages. Collectively, these findings support the hypothesis that the subcellular localization of non-apoptotic caspase cleavages may dictate substrate interactions, thereby conferring non-canonical functions to caspases in macrophage differentiation and anti-inflammatory polarization.

We also established that the non-canonical activation of caspase-8 occurs independently of other caspases. To further elucidate the mechanisms involved, we investigated whether cathepsins might contribute to this activation. Given previous evidence that CTSD can activate caspase-8 during apoptosis^43^, we assessed cathepsin activity during differentiation and polarization. We observed an increased CTSB+L activity during both processes, consistent with earlier reports that increased cathepsin activities was involved in NPM cleavage during macrophagic differentiation^39,50^. Notably, we established that the non-canonical activation of caspase-8 relies specifically on CTSB. Both pharmacological or genetic inhibition of CTSB reduced caspase-8 activation and downstream non-apoptotic functions. Although the precise mechanisms of CTSB activation in response to CSF-1 remain unclear, CTSB is a well-established effector of autophagy^51^, a process essential for macrophagic differentiation^20^. This link is further supported by the identification of Beclin-1, a key autophagy regulator^52^, as a non-apoptotic caspase substrate (data not shown). Therefore, an extensive characterization of the autophagic process during anti-inflammatory polarization is needed to better understand its role and how it can interact with non-apoptotic caspase signaling.

Mechanistically, we show that active CTSB cleaves recombinant caspase-8 *in vitro,* producing a 34kDa fragment identical to that found during macrophagic differentiation and anti-inflammatory polarization. Beyond proteolytic processing, other post-translational modifications are also known to modulate caspase-8 function. Notably, Src-mediated phosphorylation of caspase-8 at tyrosine 380, known to inhibit its activation within the DISC complex^53^, was identified during both differentiation and anti-inflammatory polarization (data not shown). This phosphorylation is likely mediated by Lyn, a member of the Src family, that we also identified as a substrate of non-apoptotic caspases. Further investigations are needed to elucidate the exact role of Lyn and its cleavage in the generation of anti-inflammatory macrophages.

Finally, we demonstrate that the inhibition of CTSB and non-apoptotic caspases, particularly caspase-8 but also caspase-7 and caspase-3, disrupts anti-inflammatory polarization and reprograms anti-inflammatory macrophages towards a pro-inflammatory phenotype. This phenotypic switch is supported by transcriptomic profiling, which reveals heightened inflammatory genes expression following CTSB or caspase-8 inhibition. Nevertheless, this analysis also revealed some distinct roles for CTSB and caspase-8 in macrophage polarization. Indeed, CTSB inhibition enhanced lymphocyte proliferation and leukocyte migration whereas caspase-8 inhibition promoted anti-viral and innate immune responses. These results underscore the therapeutic potential of targeting the CTSB-caspase-8 axis to reprogram deleterious anti-inflammatory macrophages in pathological conditions such as fibrosis and cancer. While several clinical trials have investigated caspase inhibitors in such settings, outcomes have been largely disappointing due to high toxicity and / or limited efficacy, likely resulting from the lack of specificity of pan-caspase inhibitors. For example, the pan-caspase inhibitor Emricasan, failed to reduce inflammation and fibrosis in non-alcoholic steatohepatitis (NASH) and even worsened pathology, despite being well-tolerated^54^. To overcome these challenges, we are currently developing highly specific inhibitors of non-apoptotic caspases, inspired by cleavage motifs identified during monocyte-to-macrophage differentiation. NKFD and KWFD-based molecules are now being evaluated *ex vivo* and hold promising therapeutic potential.

## Acknowledgements

This research was supported by INSERM, Côte d’Azur University, Ligue Nationale Contre le Cancer (E.K. thesis funding), foundation ARC (team label 2022-2025), INCa_19428 (PLBIO24-195), Cancéropôle PACA (Prematuration 2024), Région PACA (C.D. thesis funding) and the clinical hematology department at CHU Nice. P.A., A.J. and G.R. are members of the OPALE Carnot institute (C3M UMR-1065). We acknowledge the Etablissement Français du Sang (EFS) for providing healthy human peripheral blood, the C3M facilities (imagery, cytometry, genomic) and the CHU Nice research platform for access to Ella Automated Immunoassay System.

## Author contributions

EK and PC designed, performed the experimental work and analyzed the results. EK concepted the figures. CD, SB, MB, and MF contributed to some experiments. ND and JB generated and analyzed transcriptomic data respectively. CF, AR, JC, TC, MC, and GR participated in helpful discussions. EK wrote the manuscript. PA, JFP and AJ edited the manuscript. AJ directed the work.

## Additional information

**Correspondence** and materials requests should be addressed to Emeline Kerreneur or Dr. Arnaud Jacquel.

**Supplementary information** is available on Cellular & Molecular Immunology’s website.

## Supplementary Materials and Methods

### L-lactate secretion assay

100µL of 24H macrophages supernatants and control medium of culture is loaded in triplicates into 96-well plate. L-lactate secretion is evaluated on a YSI 2950D Biochemistry Analyzer by the difference between 24 vs 0H samples further rationalized to the number of secreting cells evaluated by flow cytometry.

### Phagocytosis

Control macrophages are pre-treated with cytochalasin D (1µg/mL) for 30 minutes at 37°C and macrophages are then incubated with FITC-beads coated with E. Coli fragments (15µg/mL) for 30 minutes at 37°C (Vybrant^TM^ Phagocytosis Assay kit, Invitrogen V6694). Macrophage phagocytic activity is then determined by flow cytometry by subtracting the mean fluorescence of macrophages and their control, diluted in trypan blue to eliminate background signals.

## Supplementary Figure legends

**Supplementary Fig. 1.**
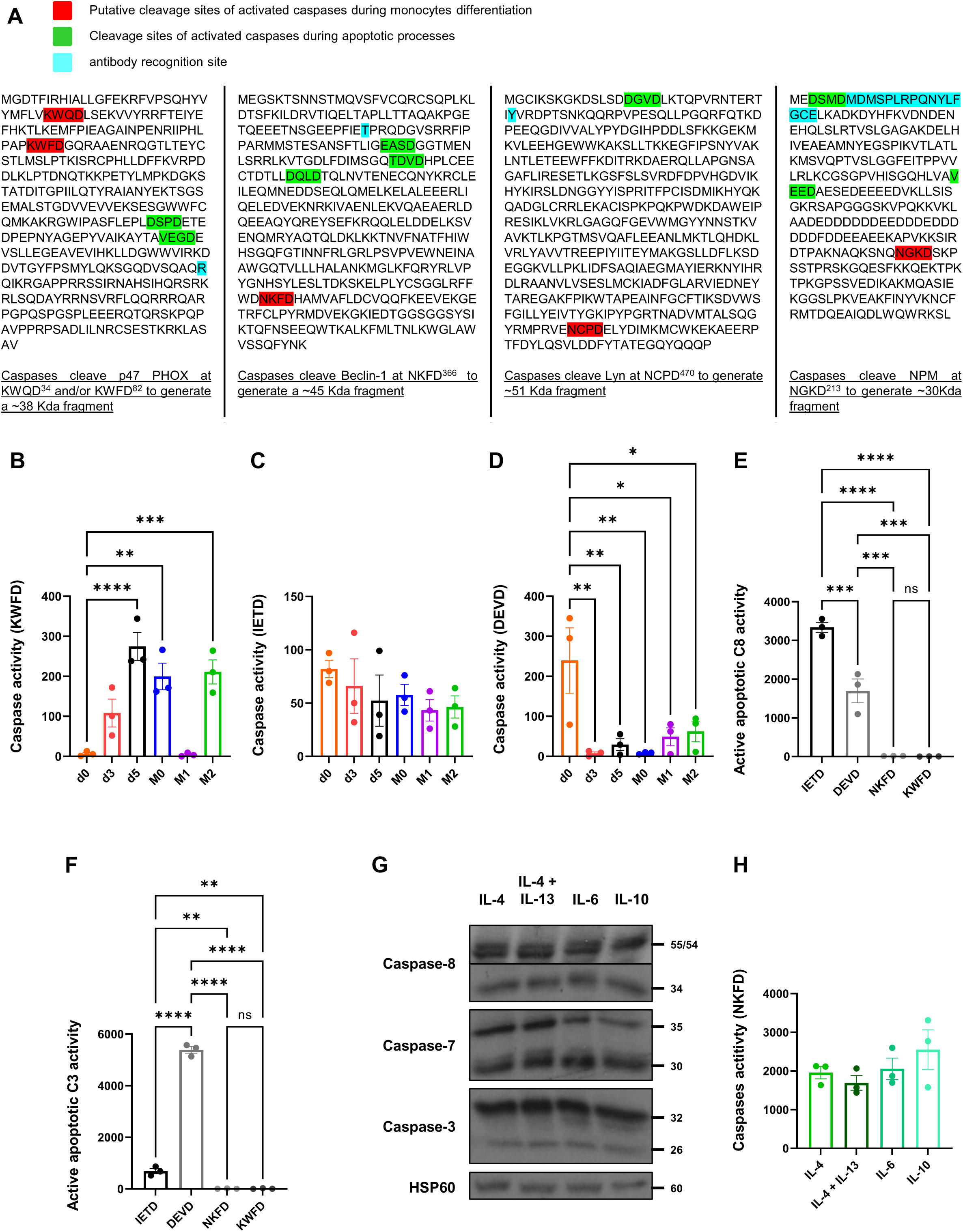
Caspase activation during monocyte-to-macrophage differentiation and anti-inflammatory macrophages polarization. **a** Amino acid sequences of caspase substrates and their putative cleavage sites during human monocyte-to-macrophage differentiation. Human monocytes (d0) are differentiated with CSF-1 for 5 days and then treated with CSF-1 (M0) or polarized with IFN-γ + LPS for pro-inflammatory macrophages, IL-4 (M2), IL-4 ± IL-13, IL-6 or IL-10 for anti-inflammatory macrophages for 2 days. **b-d** Enzymatic measurement of non-apoptotic caspase activity in differentiating and polarized macrophages lysates using a specific fluorescent peptide (**b:** Ac-KWFD-AMC; **c:** Ac-IETD-AMC; **d:** Ac-DEVD-AMC). Results are expressed as A.U/min/mg of proteins and represent the mean ± SEM of 3 independent experiments performed in triplicates. **e-f** Enzymatic measurement of recombinant caspase-8 (**e**) or caspase-3 (**f**) activity using specific fluorescent peptides (Ac-IETD-AMC, Ac-DEVD-AMC, Ac-NKFD-AMC and Ac-KWFD-AMC). Results are expressed as A.U/min/mg of proteins and represent the mean ± SEM of 3 independent experiments performed in triplicates. **g** Caspase-8, caspase-7 and caspase-3 expression analysis by immunoblotting. HSP60 is used as a loading control. **h** Enzymatic measurement of non-apoptotic caspase activity using a specific fluorescent peptide (Ac-NKFD-AMC). Results are expressed as A.U/min/mg of proteins and represent the mean ± SEM of 3 independent experiments performed in triplicates. ns : p > 0.05 ; * p < 0.05 ; ** p < 0.01 ; *** p < 0.001 and **** p < 0.0001 according to an ordinary one-way ANOVA.

**Supplementary Fig. 2.**
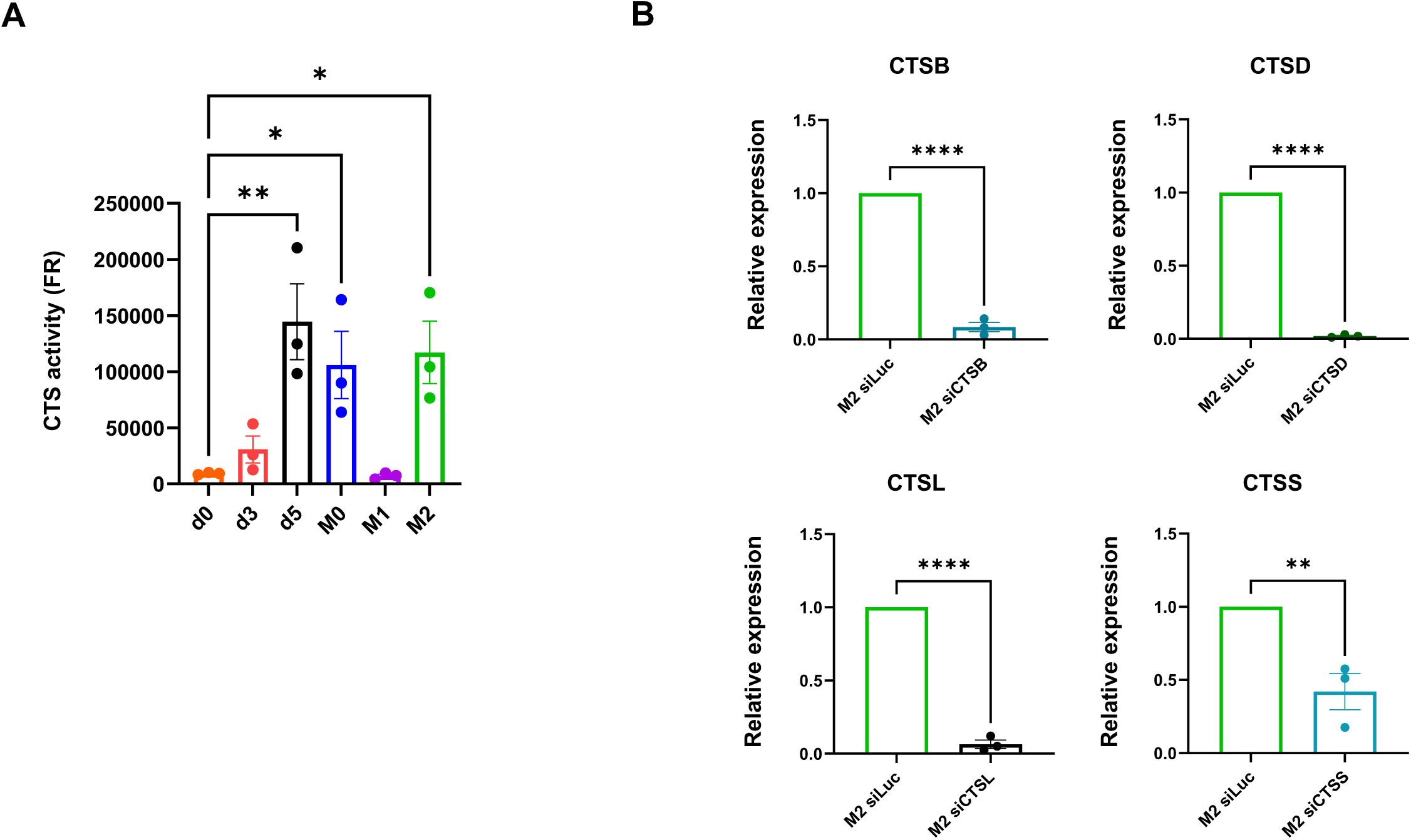
CTS enzymatic activity and expression in polarizing macrophages. **a** Human monocytes (d0) are differentiated with CSF-1 for 5 days and then treated with CSF-1 (M0) or polarized with IFN-γ + LPS (M1) or IL-4 (M2) for 2 days. Enzymatic measurement of CTS using a specific fluorescent peptide Ac-FR-AMC. Results are expressed as A.U/min/mg of proteins and represent the mean ± SEM of 3 independent experiments performed in triplicates. * p < 0.05 and ** p < 0.01 according to an ordinary one-way ANOVA. **b** Anti-inflammatory polarizing macrophages are transfected with siRNA directed against Luciferase, CTSB, D, L, or S for 2 days. RT-qPCR analysis of CTSB, D, L and S gene expression. Results are expressed as relative expression and represent the mean ± SEM of 3 independent experiments. ** p < 0.01 and **** p < 0.0001 according to a two-tailed unpaired Student’s t test.

**Supplementary Fig. 3.**
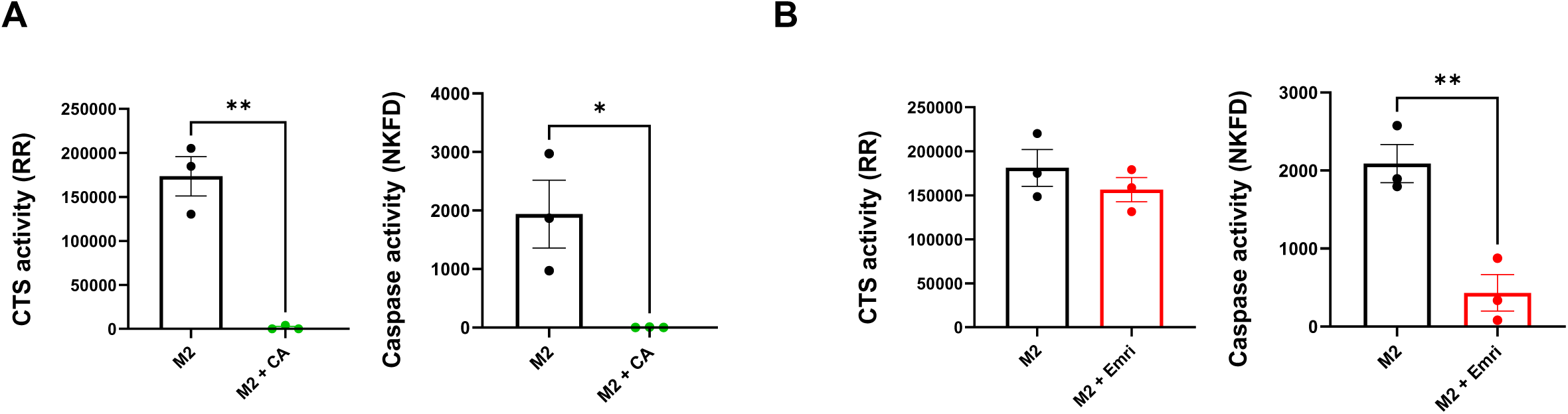
CTS and non-apoptotic caspase activities in anti-inflammatory macrophages upon pharmacological CTSB or caspase inhibition. Enzymatic measurement of CTSB+L and non-apoptotic caspase activities using a specific fluorescent peptide (Ac-RR-AMC and Ac-NKFD-AMC respectively). Results are expressed as A.U/min/mg of proteins and represent the mean ± SEM of 3 independent experiments performed in triplicates. ns : p > 0.05 ; * p < 0.05 and ** p < 0.01 according to a two-tailed unpaired Student’s t test.

**Supplementary Fig. 4.**
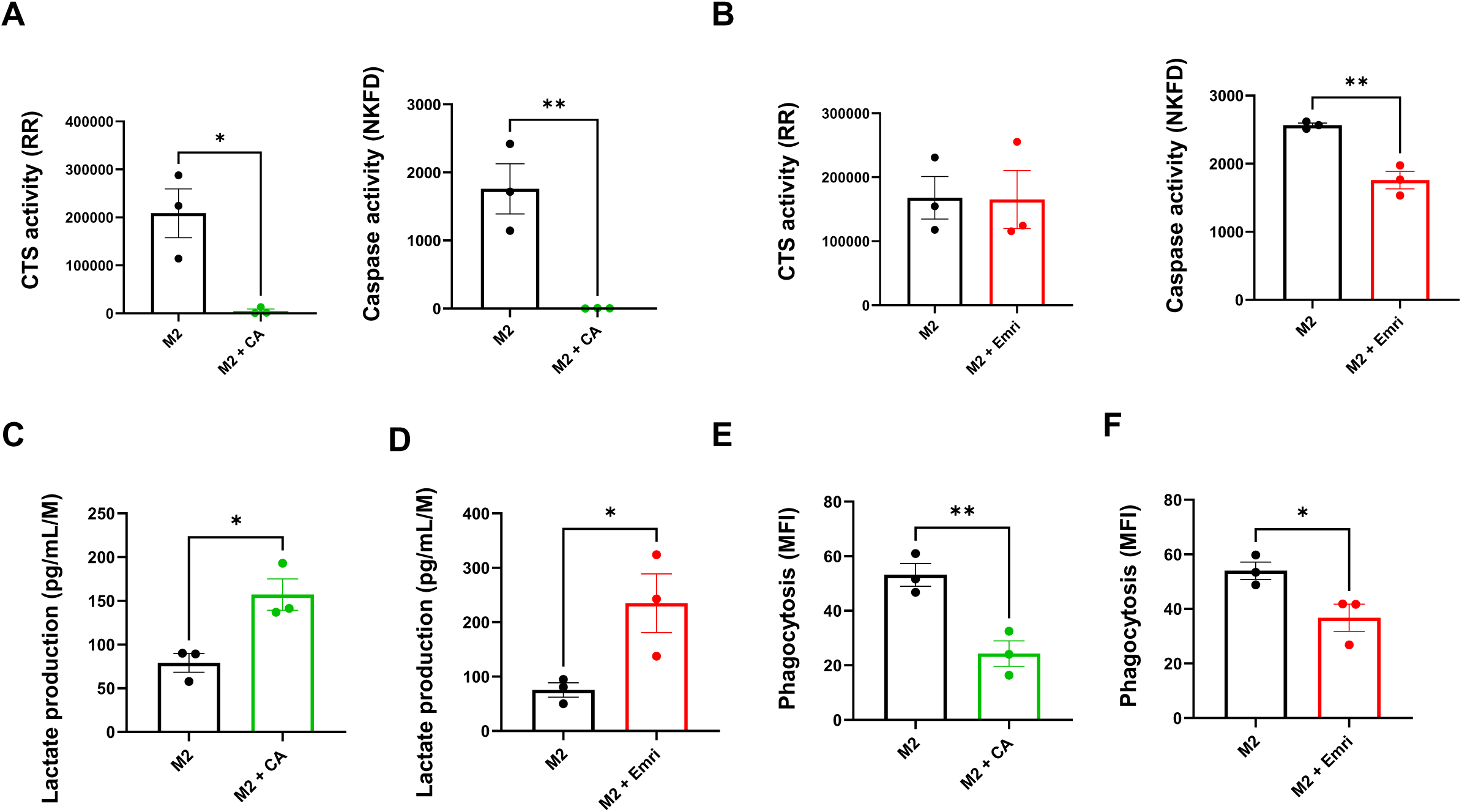
Phenotype and functional profile of anti-inflammatory macrophages upon CTSB or caspase pharmacological inhibition. Human monocytes are differentiated in macrophages with CSF-1 for 5 days and polarized into anti-inflammatory macrophages during 2 days with IL-4 and further treated with Emricasan or CA-074 for 2 days. **a-b** Enzymatic measurement of CTSB and non-apoptotic caspase activities using specific fluorescent peptides (Ac-RR-AMC and Ac-NKFD-AMC respectively). Results are expressed as A.U/min/mg of proteins and represent the mean ± SEM of 3 independent experiments performed in triplicates. **c-d** Lactate production analysis. Results are expressed as pg/mL/10^6^ cells and represent the mean ± SEM of 3 independent experiments. **e-f** Phagocytic capacity analysis of macrophages by flow cytometry. Results are expressed as MFI (Mean Fluorescence Intensity) and represent the mean ± SEM of 3 independent experiments. ns : p > 0.05 ; * p < 0.05 and ** p < 0.01 according to a two-tailed unpaired Student’s t test.

**Supplementary Fig. 5.**
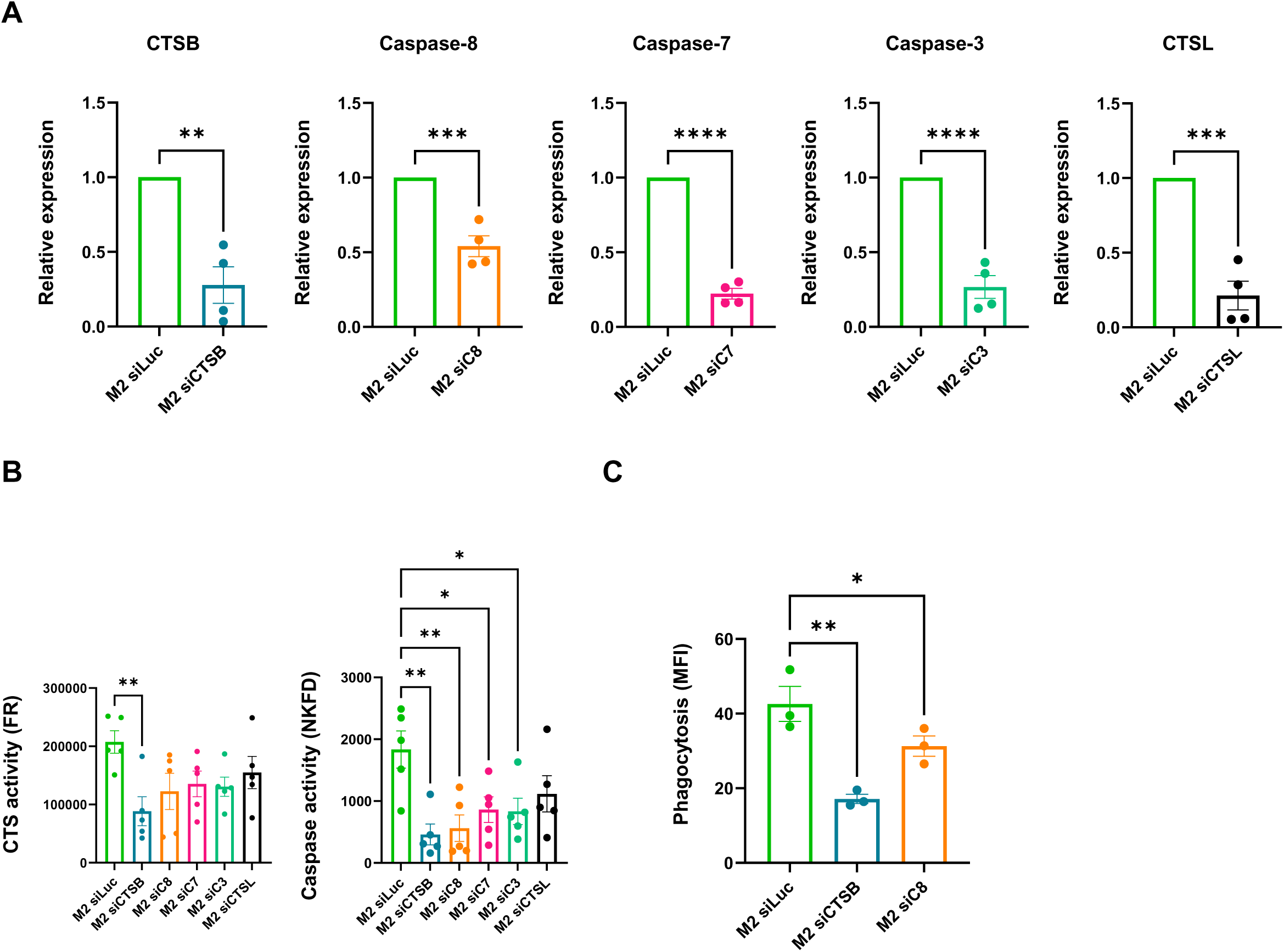
Phenotype and functional profile of anti-inflammatory macrophages upon genetic inhibition of CTSB, caspase-8, caspase-7, caspase-3 or CTSL. Anti-inflammatory macrophages are transfected with siRNA directed against Luciferase (used as a control of transfection), caspase-8, caspase-7, caspase-3, CTSB or CTSL during 2 (**a**) or 3 days (**b, c**). **a** RT-qPCR analysis of CTSB, caspase-8, caspase-7, caspase-3 and CTSL gene expression. Results are expressed as relative expression and represent the mean ± SEM of 4 independent experiments. ** p < 0.01 ; *** p < 0.001 and **** p < 0.0001 according to a two-tailed unpaired Student’s t test. **b** Enzymatic measurement of CTSB and non-apoptotic caspase activities using specific fluorescent peptides (Ac-FR-AMC and Ac-NKFD-AMC respectively). Results are expressed as A.U/min/mg of proteins and represent the mean ± SEM of 5 independent experiments performed in triplicates. **c** Phagocytic capacity analysis by flow cytometry. Results are expressed as MFI (Mean Fluorescence Intensity) and represent the mean ± SEM of 3 independent experiments. ns : p > 0.05 ; * p < 0.05 ; ** p < 0.01 and **** p < 0.0001 according to an ordinary one-way ANOVA.

**Supplementary Fig. 6.**
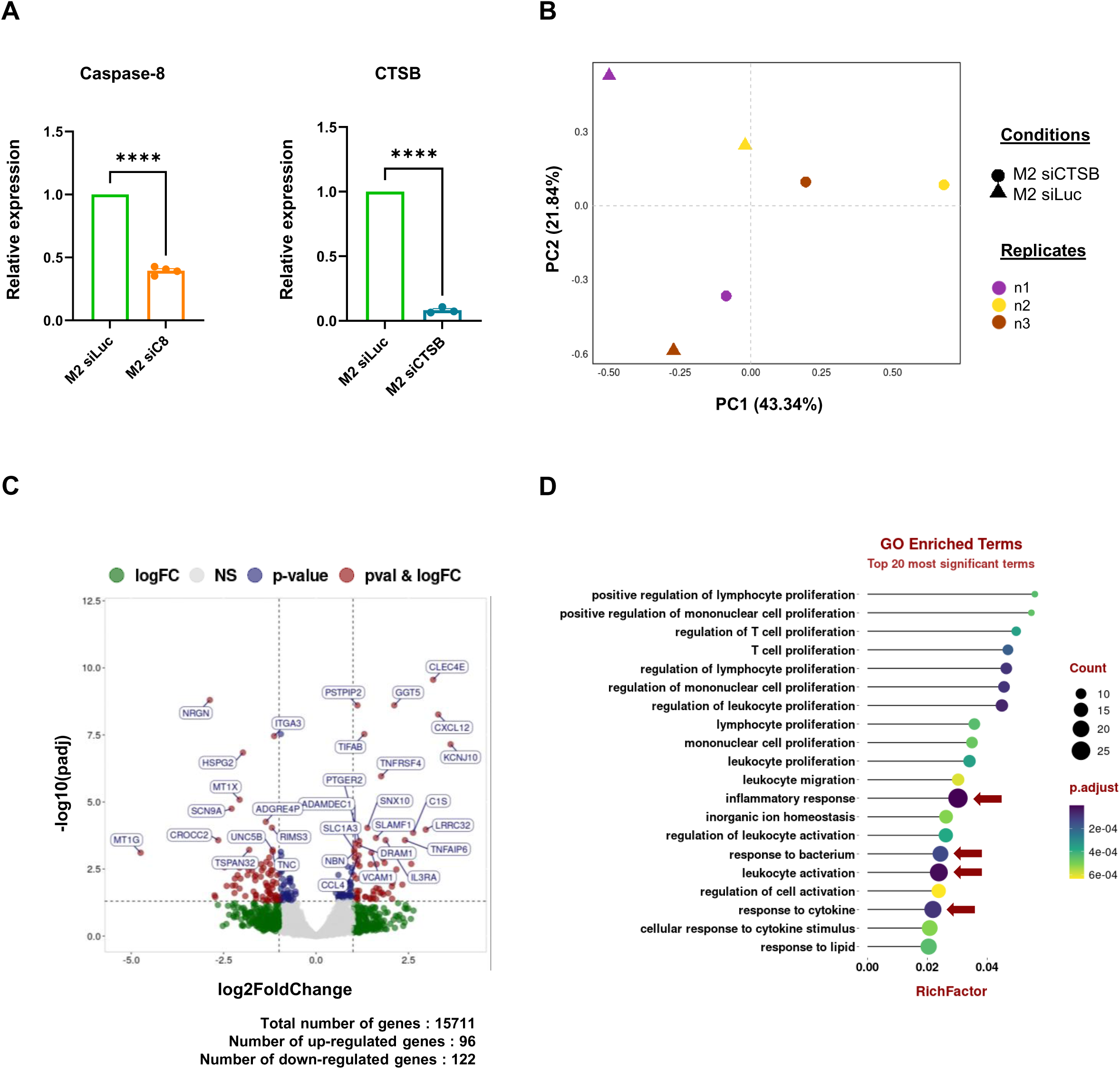
Transcriptomic reprogramming of anti-inflammatory macrophages under genetic inhibition of CTSB. Anti-inflammatory macrophages are transfected with siRNA directed against Luciferase (M2 siLuc), used as a control of transfection, caspase-8 (M2 siC8) or CTSB (M2 siCTSB) for 2 days. **a** RT-qPCR analysis of caspase-8 or CTSB gene expression. Results are expressed as relative expression and represent the mean ± SEM of 4 or 3 independent experiments respectively. ns : p > 0.05 ; * p < 0.05 ; ** p < 0.01 and *** p < 0.001 according to according to a two-tailed unpaired Student’s t test. **b** Unsupervised 2D projection of the six transcriptome samples principal component analysis. **c** Volcano plot representation of significant dysregulated genes between M2 siCTSB compared to M2 siLuc. **d** Dot plot representation of the top 20 biological processes from gene ontology analysis.

